# Fast computation of principal components of genomic similarity matrices

**DOI:** 10.1101/2022.10.06.511168

**Authors:** Georg Hahn, Sharon M. Lutz, Julian Hecker, Dmitry Prokopenko, Michael H. Cho, Edwin K. Silverman, Scott T. Weiss, Christoph Lange

## Abstract

The computation of a similarity measure for genomic data, for instance using the (genomic) covariance matrix, the Jaccard matrix, or the genomic relationship matrix (GRM), is a standard tool in computational genetics. The principal components of such matrices are routinely used to correct for biases in, for instance, linear regressions. However, the calculation of both a similarity matrix and its singular value decomposition (SVD) are computationally intensive. The contribution of this article is threefold. First, we demonstrate that the calculation of three matrices (the genomic covariance matrix, the weighted Jaccard matrix, and the genomic relationship matrix) can be reformulated in a unified way which allows for an exact, faster SVD computation. An exception is the Jaccard matrix, which does not have a structure applicable for the fast SVD computation. An exact algorithm is proposed to compute the principal components of the genomic covariance, weighted Jaccard, and genomic relationship matrices. The algorithm is adapted from an existing randomized SVD algorithm and ensures that all computations are carried out in sparse matrix algebra. Second, an approximate Jaccard matrix is introduced to which the fast SVD computation is applicable. Third, we establish guaranteed theoretical bounds on the distance (in *L*_2_ norm and angle) between the principal components of the Jaccard matrix and the ones of our proposed approximation, thus putting the proposed Jaccard approximation on a solid mathematical foundation. We illustrate all computations on both simulated data and data of the 1000 Genome Project, showing that the approximation error is very low in practice.

## 1 Introduction

In computational genomics, the computation of eigenvectors as part of a principal component analysis (PCA) is a widespread method to infer population structure and to correct for confounding due to ancestry. An extensive amount of publications in the literature are devoted to this topic.

It has long been known that case-control studies are subject to population stratification which can induce significant spurious associations at loci that are unrelated with a response (Reich and Goldstein, 2001). For instance, Campbell et al. (2005) showcase such a spurious association through stratification by showing that a SNP in the lactase gene LCT varies widely in frequency across Europe and was strongly associated with height. To correct for stratification in genome-wide association studies (Patterson et al., 2006), methodology such as EIGENSTRAT (Price et al., 2006) has been subsequently developed. Further computational improvements are available (Li and Yu, 2008). Further works in the literature address confounding induced by population stratification with the help of a two-step procedure (Epstein et al., 2007), or PCA analyses with tens of thousands of single-nucleotide polymorphisms (SNPs) to infer population structure (Lee et al., 2012).

This article focuses on the fast computation of eigenvectors of four different similarity matrices employed in computational genomics. These matrices provide pairwise similarity measures between genomes, and their eigenvectors are popular means to correct for population stratitification. We consider the (classic) genetic covariance matrix (Schäfer and Strimmer, 2005), the Jaccard matrix (Jaccard, 1901; Prokopenko et al., 2016), the weighted Jaccard matrix (Schlauch et al., 2017), and the genomic relationship matrix (GRM) (Yang et al., 2011). All four matrices are computed on an input matrix 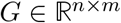, where 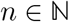 is the number of loci and 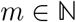 is the number of individuals. The genomic input data *G* is usually a sparse matrix. All four matrices result in an output matrix of dimension *m × m*, where each entry (*i,j*) is a similarity measure between the genomic data of individuals *i* and *j*. All four matrices are symmetric by definition.

In real data applications such as the 1000 Genomes Project (The 1000 Genomes Project Consortium, 2015) or the UK Biobank (Sudlow et al., 2015), the number of individuals quickly reaches numbers in the thousands, ten thousands, or hundred thousands. In this case, traditional eigenvector computations, for instance using the function *eigen* in R (R Core Team, 2014), become infeasible. Alternatives with lower computational complexity are iterative methods such as the *power method* (von Mises and Pollaczek-Geiringer, 1929), also called Von Mises iteration, or the truncated singular value decomposition (SVD) implemented in, for instance, the R-package *RSpectra* (Qiu et al., 2022). However, for these alternatives to be applicable, the complete similarity matrix has to be computed first. In the best case, calculating each similarity matrix is computationally intensive itself, while in the worst case, its calculation is computationally infeasible as the genomic input *X* is usually sparse, whereas each of the four aforementioned similarly measures are typically dense. As shown in this work, the computation of the similarity measure can be omitted entirely for the computation of its eigenvectors.

The contribution of this article is threefold. First, we show that the eigenvectors of three similiary matrices (the genetic covariance matrix, the weighted Jaccard matrix, and the genomic relationship matrix) can be computed efficiently by rewriting their computations in a unified way which allows for an exact, faster computation. We then propose a tailored algorithm by adapting an existing randomized SVD algorithm of Halko et al. (2011). Our algorithm never actually computes a similarity matrix and fully supports sparse matrix algebra for efficient calculations. However, we will show that the same trick does not apply to the Jaccard matrix. Second, we therefore propose an approximate Jaccard matrix which likewise allows for an efficient computation of its eigenvectors without actually computing the similarity measure. Third, we establish guaranteed theoretical bounds on the distance (in *L*_2_ norm and angle) between the eigenvectors of the Jaccard matrix and the ones of our proposed approximation, thus putting the proposed Jaccard approximation on a solid mathematical foundation.

The computation of a randomized PCA has also been considered in Abraham and Inouye (2014). In their publication, the authors likewise adapt the original algorithm of Halko et al. (2011). However, the authors only consider the genomic relationship matrix and do not establish the unified framework which actually allows for the PCA computation of the other similarity matrices as well. Moreover, the Jaccard approximation and the theoretical bounds we prove are unconsidered.

In an experimental section, we illustrate the proven exactness of the proposed computations for the genetic covariance matrix, the weighted Jaccard matrix, and the genomic relationship matrix.

Special attention is given to the Jaccard matrix. Using simulated data, we verify the proven theoretical bounds on the distance between the eigenvectors of the Jaccard matrix and our proposed approximation, showing that the approximation error is very low in practice. Moreover, we demonstrate the trade-off in accuracy between the Jaccard matrix and its approximation for population stratification plots using data of the 1000 Genomes Project.

The paper is structured as follows. Section 2 introduces the proposed decomposition of the four similarity matrices under consideration (Section 2.1), establishes that the fast computation of eigenvectors applies to three of these matrices (Section 2.2), introduces a new approximation of the Jaccard similarity matrix (Section 2.3), establishes theoretical bounds on the accuracy of the approximation (Section 2.4), and summarizes all findings as an efficient algorithm (Section 2.5) together with asymptotic runtime considerations (Section 2.6). All experimental results can be found in Section 3. The article concludes with a discussion in Section 4.

The proposed methodology has been implemented as part of the R-package *locStra* (Hahn et al., 2021, 2022), available on the Comprehensive R Archive Network (R Core Team, 2014). In the entire article, 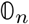 and 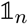 denote the vectors of length *n* with all entries set to 0 or 1, respectively. Moreover, we denote with *σ_r_*(*Y*) and *σ_c_*(*Y*) the row and column sums of a matrix *Y*, and with *μ_r_*(*Y*) and *μ_c_*(*Y*) the row and column means of *Y*, respectively. Finally, the notation *v* ⊗ *w* is used to denote the outer product between two vectors *v* and *w*.

## 2 Methods

This section demonstrates that the genetic covariance matrix, the weighted Jaccard matrix, and the genomic relationship matrix can be expressed in a unified way which allows for an efficient computation of their eigenvectors (Section 2.1 and Section 2.2). This does not apply to the Jaccard matrix, for which we propose a new approximation instead that allows for a fast eigenvector computation (Section 2.3). Importantly, we establish theoretical bounds on the accuracy of the eigenvectors obtained from the Jaccard approximation (Section 2.4). We summarize all findings in an algorithm tailored to the four similarity matrices in Section 2.5. The section concludes with considerations on the asymptotic speedup in Section 2.6.

### 2.1 Fast computation of eigenvectors

The algorithm of Halko et al. (2011) allows one to compute the eigenvectors of either the matrix *X*^⊤^*X*, or the matrix *XX*^⊤^, by considering *X* only. The actual product *X*^⊤^*X* or *XX*^⊤^ does not need to be computed at any point in time. This is especially advantageous if the matrix *X* is sparse since then, oftentimes, *X*^⊤^*X* and *XX*^⊤^ are dense. In the following, we focus on the computation of the eigenvectors of *X*^⊤^*X* only.

In order to apply the fast eigenvector computation of Halko et al. (2011), we need to express all similarity matrices under consideration as a product of the form *X*^⊤^*X*. This is not a straightforward task, as the computation of the aforementioned similarity matrices involves normalization and centering operations. As a first step, we consider the decompositions

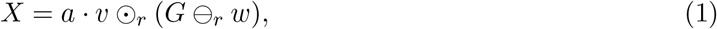

or alternatively, as

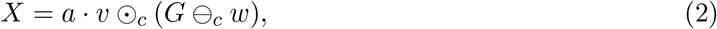

where 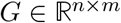 is the genomic input data on 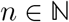 loci and 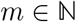 individuals, 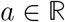 is a scalar, and *v, w* are vectors of appropriate dimensions. The scalar *a* is kept separate and not absorbed into *v* for clarity of notation, as most similiarity matrices have a separate normalizing constant. The operations ʘ*_r_*, ⊖*_r_* as well as ʘ*_c_*, ⊖*_c_* are the row/column-wise multiplication and subtraction operation of a vector with a matrix, respectively. To be precise, *G* ⊖*_r_ w* subtracts *w* from all rows of *G*, and *G* ⊖*_c_ w* subtracts *w* from all columns of *G*. Analogously, *v* ʘ_r_ *G* multiplies all rows of *G* with *v*, and *v* ⊖_c_ *G* multiplies all columns of *G* with *v* (assuming *v* and *w* are of appropriate dimensions).

The specific decomposition in the form *X*^⊤^*X* using eq. (1) and eq. (2) has two crucial advantages:

1. As shown in the following sections (Section 2.2 and Section 2.3), the genetic covariance matrix, the weighted Jaccard matrix, the genomic relationship matrix, and a newly proposed Jaccard approximation can be expressed in a unified form as *X*^⊤^*X*, with *X* as in eq. (1) and eq. (2).
2. While *G* is usually a sparse matrix, centering *G* with a vector *w* as done in eq. (1) and eq. (2) usually results in a dense matrix, which is computationally inefficient to handle or even infeasible. As shown in Section 2.5, the main advantage of eq. (1) and eq. (2) consists in the fact that they allow one to compute eigenvectors in sparse algebra only, without ever performing the multiplication or subtraction operations.

### 2.2 Decomposition of three similarity matrices

The genetic covariance matrix, the weighted Jaccard matrix, and the genomic relationship matrix allow for an expression of the form of eq. (1) or eq. (2):

1. The genetic covariance matrix is computed as 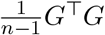 after centering all rows of *G* with their respective column means. This fits into the framework of eq. (1) by setting 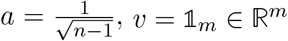, and 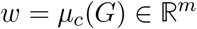.
2. The computation of the weighted Jaccard matrix of Schlauch et al. (2017) is more involved and repeated here for convenience. First, a quantity *numAlleles* is computed as 2*n*. Then, the sum of variants in *G* is computed as the row sums of *G* and denoted as *sumVariants.* In a pre-processing step to invert the minor alleles, all rows in *G* are inverted if their sum of variants is strictly larger than *n.* Second, a set of weights is computed as follows. A quantity *totalPairs* is computed as *s*(*s* – 1)/2, where 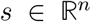 is the vector of row sums of *G* and the vector multiplication is performed componentwise. The weight vector *weights* is then computed as *numAlleles*\*(*numAlleles*–1)/*totalPairs,* again taking all operations to be componentwise. The vectors *totalPairs* and *weights* both have dimension *n*. Third, the weighted Jaccard matrix is computed as 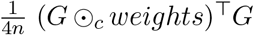. This computation fits into the framework of eq. (2) by setting 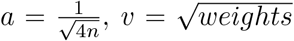 (with the square root operation performed componentwise), and 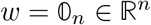.
3. The genomic relationship matrix (GRM) of Yang et al. (2011) exists in two flavors, a robust and a non-robust version. Both are easily defined as follows. Define *p* = *μ_r_* (*G*)/2 as row means of *G*, and *q* = 2*p*(1 – *p*), where the vector multiplication is again understood componentwise. Let *s* be the sum of all entries in *q*. After centering the columns of *G* with 2*p* (that is, *X* ← *X* ⊖*_c_* (2*p*)), the robust GRM is defined as *G*^⊤^*G/s*, and the non-robust GRM is defined as 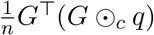. Both the robust and non-robust versions of the GRM fit into the framework of eq. (2). For the robust GRM, we set 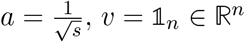, and *w* = 2*p*. For the non-robust GRM, we set 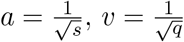, again taking all vector operations to be componentwise, and *w* = 2*p*.

Finally, it remains to note that the Jaccard matrix (Prokopenko et al., 2016) does not allow for a decomposition into *X*^⊤^*X* with an appropriately chosen matrix *X*. This is easily verified in practice. Indeed, it is not complicated to find a simulated or real life genomic dataset for which the Jaccard matrix has negative eigenvalues, and is thus not positive (semi-)definite. This proves that a decomposition into the form *X*^⊤^*X*, which necessarily implies positive (semi-)definiteness, is impossible.

### 2.3 A new approximation of the Jaccard similarity matrix

The Jaccard similarity matrix (Prokopenko et al., 2016) is computed as follows on a *binary* genomic input matrix *G*. First, a matrix 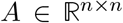 is computed. Each entry (*i, j*) in *A* is obtained by computing the logical *and* operation on the binary columns *i* and *j* of *G*, and storing the sum of ones (or values *True*) in the resulting vector in *A_i,j_*. Similarly, a matrix 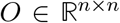 is computed whose entry (*i,j*) represents the sum of ones after an or operation on the binary columns *i* and *j* of *G*. The Jaccard matrix *J* is then computed as *J* = *A/O,* where the matrix division is taken componentwise.

It is important to note that for binary matrices, the logical *and* operation required to compute 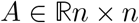 is equivalent to simply computing the matrix-matrix product of *G* with its transpose, that is *A* = *G*^⊤^*G*. Therefore, it is in fact the *or* operation that prevents the Jaccard matrix from being expressible in the form *X*^⊤^*X* as shown in Section 2.1.

To fix this, we propose a simple approximation that replaces the computation of the matrix *O*. Note that, when computing the logical *or* operation on two columns *i* and *j* of *G*, the maximal number of ones we can obtain in the resulting vector is 2 max(s), where *s* is the vector of column sums of *G*. This case occurs if both columns of *G* contain the maximal number of nonzero entries, though at different positions each. To be conservative, we therefore propose to compute an approximation of *J* = *A/O* as 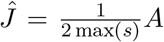. The matrix 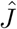 has the property that any entry in 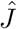 is at most as large as the corresponding one in *J*, thus never overestimating the similarity between two individuals.

Due to its simpler structure, the approximate Jaccard matrix 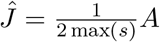 fits into the frame-work of eq. (1) by taking 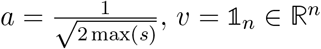, and 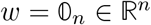, where *s* was the vector of column sums of *G*.

### 2.4 Theoretical error bounds on the eigenvectors of the Jaccard approximation

The original Jaccard matrix as defined in Prokopenko et al. (2016) and the proposed approximate Jaccard matrix of Section 2.3 naturally differ slightly, and so do their eigenvectors.

Therefore, our proposed approximate Jaccard matrix comes with a classical speed/ accuracy tradeoff. It is much faster to compute, though at the expense of a slight loss in accuracy. However, as the computation of eigenvectors is at the heart of this paper, this section establishes guaranteed theoretical bounds on the distance (in *L*_2_ norm and angle) between the eigenvectors of the Jaccard matrix and the ones of our proposed approximation. These a priori bounds allow the user to gauge in advance the trade-off between obtained speedup and sacrificed accuracy.

To derive the bounds, we make use of the so-called “Davis-Kahan sin(θ)” theorem (Davis and Kahan, 1970). Citing the statement of the theorem in Rigollet (2020), let 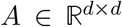 and 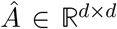 be two symmetric matrices with eigen decompostions given by 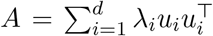 and 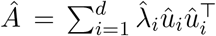, where *λ*_1_ ≥ … *λ_d_* and 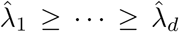 are the sorted eigenvalues. Then, 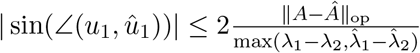 and 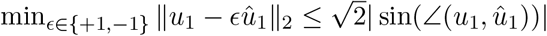.

The Davis-Kahan theorem therefore allows one to bound the distance (up to multiplication with +1 or −1, since eigenvectors are only defined up to a unit) between the eigenvectors of a matrix *A* and an arbitrary perturbation 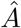 of *A* using the angle between their eigenvectors, which in turn is bounded by a quantity involving the operator norm of 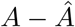 and their first eigenvalues.

Applied to the Jaccard matrix *J* and our proposed approximation 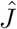 of Section 2.3, we see that the difference between the eigenvectors *u*_1_ of *J* and 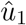 of 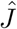, up to a unit {+1, −1}, can be bounded

#### Algorithm 1

randomized sparse SVD for *X*^⊤^*X*, where *X* = *a* · *v* ʘ*_c_* (*G* ⊖*_c_ w*)

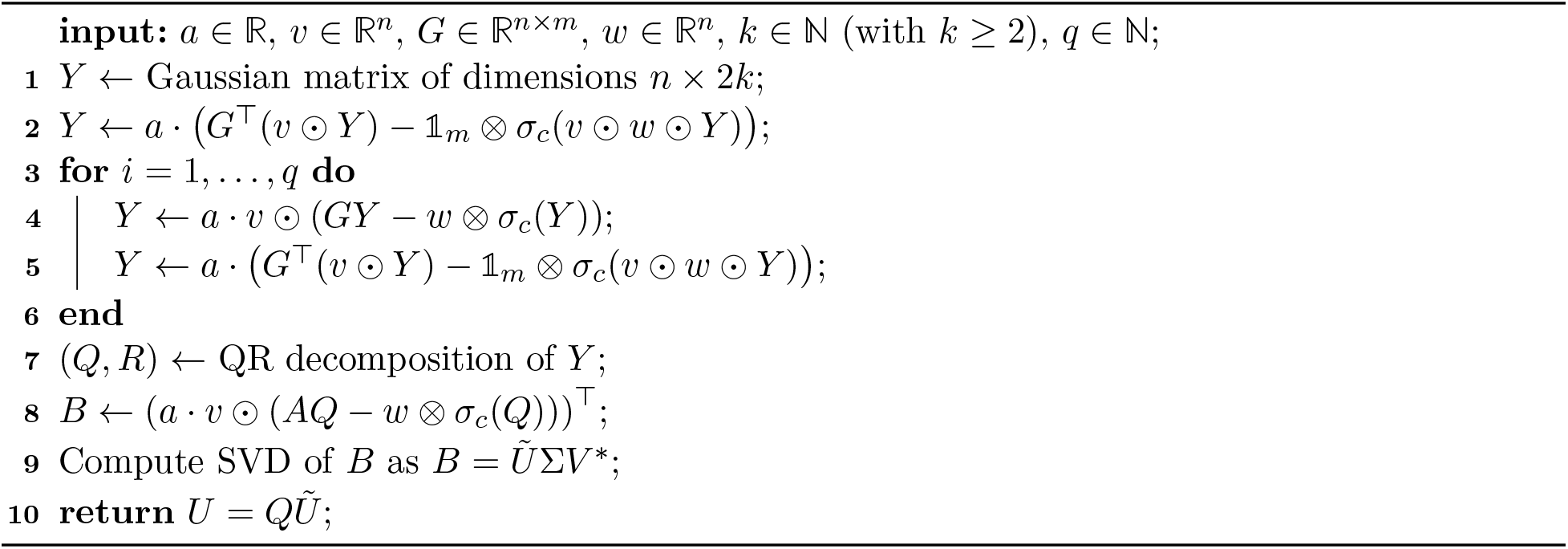

as

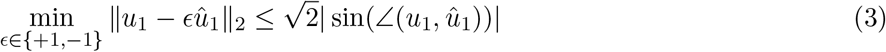

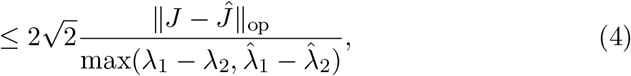

where *λ*_1_, *λ*_2_ and 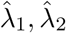 are the first two eigenvalues of *J* and 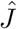, respectively. The operator norm of the two matrices in eq. (4) is straightforward to compute, and the first two eigenvalues of the two matrices can be computed efficiently using, for instance, the power method (also called von Mises iteration) of von Mises and Pollaczek-Geiringer (1929).

It is important to note that the aforementioned approximation can also be used without actually computing the eigenvalues of *J* and 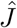. This is possible with the help of the Gershgorin circle theorem (Gerschgorin, 1931), which allows one to easily obtain lower and upper bounds on all eigenvalues of a matrix.

### 2.5 An efficient algorithm using sparse matrix algebra

The decomposition of eq. (1) and eq. (2) allows one to formulate an efficient algorithm to compute the eigenvectors of *X*^⊤^*X* in sparse algebra only, without ever performing the multiplication or subtraction operations. This method is given in Algorithm 1 for eq. (2), which is adapted from the algorithm of Halko et al. (2011) by exploiting sparse matrix operations only. An adaptation to eq. (1) is straightforward and thus omitted in this paper.

The input of Algorithm 1 is the matrix *X* = *a* · *v* ʘ*_c_* (*G* ⊖*_c_ w*) of which one wishes to compute the eigenvectors of *X*^⊤^*X*, the number of desired eigenvectors 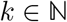, as well as an auxiliarly parameter 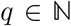 to control the numerical accuracy. Typically, the choice *q* = 2*k* is recommended in Halko et al. (2011), where larger values provide higher numerical accuracy.

In Algorithm 1, after initializing the matrix *Y* with Gaussian random numbers, the operation *Y* ← (*XX*\*)*^q^XY* is performed as done in Halko et al. (2011). As seen in lines 2-5, the structure of *X* = *a·vʘ_c_*(*G*⊖*_c_w*) allows one to separate the exponentiation into a part using sparse matrix algebra operations for *G* only, and simple outer products of lower dimension. Next, a QR decomposition is computed for *Y*, and *B* ← *Q*\**X* is computed which again can be separated into a part using sparse matrix algebra only, and an outer product of lower dimensions. The resulting matrix B has the dimensions (2*k*) × *n* only, thus allowing for a fast SVD computation. Algorithm 1 returns 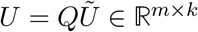 as done in Halko et al. (2011).

Note that the matrix *R* of the QR decomposition, as well as the matrices Σ and *V*\* of the SVD are actually not used in the algorithm. Their computation can therefore be omitted.

### 2.6 Runtime considerations

As shown in Hahn et al. (2021), the effort to compute the four similarity matrices is *O*(*nm*^2^). If sparse algebra is used, the effort is *O*(*snm*^2^), where *s* ∈ [0, 1] is the matrix sparsity parameter. The resulting similarity matrix has dimensions *m* × *m* in all cases. A subsequent standard SVD to compute their eigenvectors has effort *O*(*m*^3^) (Golub and Van Loan, 1996). Together, the effort for the computation of the similiary matrix and its eigenvectors via classic SVD is *O*(*nm*^2^ + *m*^3^).

Using Algorithm 1, the computation of any of the four similarity matrices is entirely omitted. Running Algorithm 1 has a computational effort of *O*(*qk* · *nm* + *k*^2^(*n* + *m*)), thus making the computation of the first 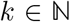 eigenvectors of any similarity matrix a linear operation in both *m* and *n*.

## 3 Experimental results

This section first presents experimental results on the numerical quality of the eigenvectors returned by Algorithm 1 when applied to the genetic covariance matrix, the weighted Jaccard matrix, and the genomic relationship matrix (Section 3.1). Afterwards, we examine the numerical accuracy of the eigenvectors computed for the proposed approximate Jaccard matrix in comparison with the original Jaccard matrix (Section 3.2), and verify the theoretical bounds derived in Section 2.4.

### 3.1 Application to the 1000 Genomes Project data

We apply Algorithm 1 to chromosome 1 of the 1000 Genomes Project (The 1000 Genomes Project Consortium, 2015), with the aim to examine the accuracy of the first two eigenvectors in population stratification figures.

We first prepare the raw data of the 1,000 Genome Project using PLINK2 (Purcell and Chang, 2019) using a cutoff value 0.01 for the option --*max-maf* to select rare variants. Moreover, we employ LD pruning with parameters --*indep-pairwise 2000 10 0.01*. All results focus on the European super population, containing 503 subjects and approximately 5 million rare variants.

Figure 1 shows results for the first two eigenvectors of the genomic relationship matrix (GRM), colored by subpopulation (GBR, FIN, IBS, CEU, TSI). Eigenvectors are once computed by constructing the GRM matrix fully on the 1000 Genomes Project dataset before calculating its eigenvectors, and once using Algorithm 1 in connection with eq. (2). We observe that the plots are visibly identical. This is to be expected, as the GRM matrix allows for an exact decomposition in the from *X*^⊤^*X* suitable for Algorithm (1), see Section 2.2. Similar plots for the genetic covariance matrix and the weighted Jaccard matrix can be found in Section A.

**Figure 1:**
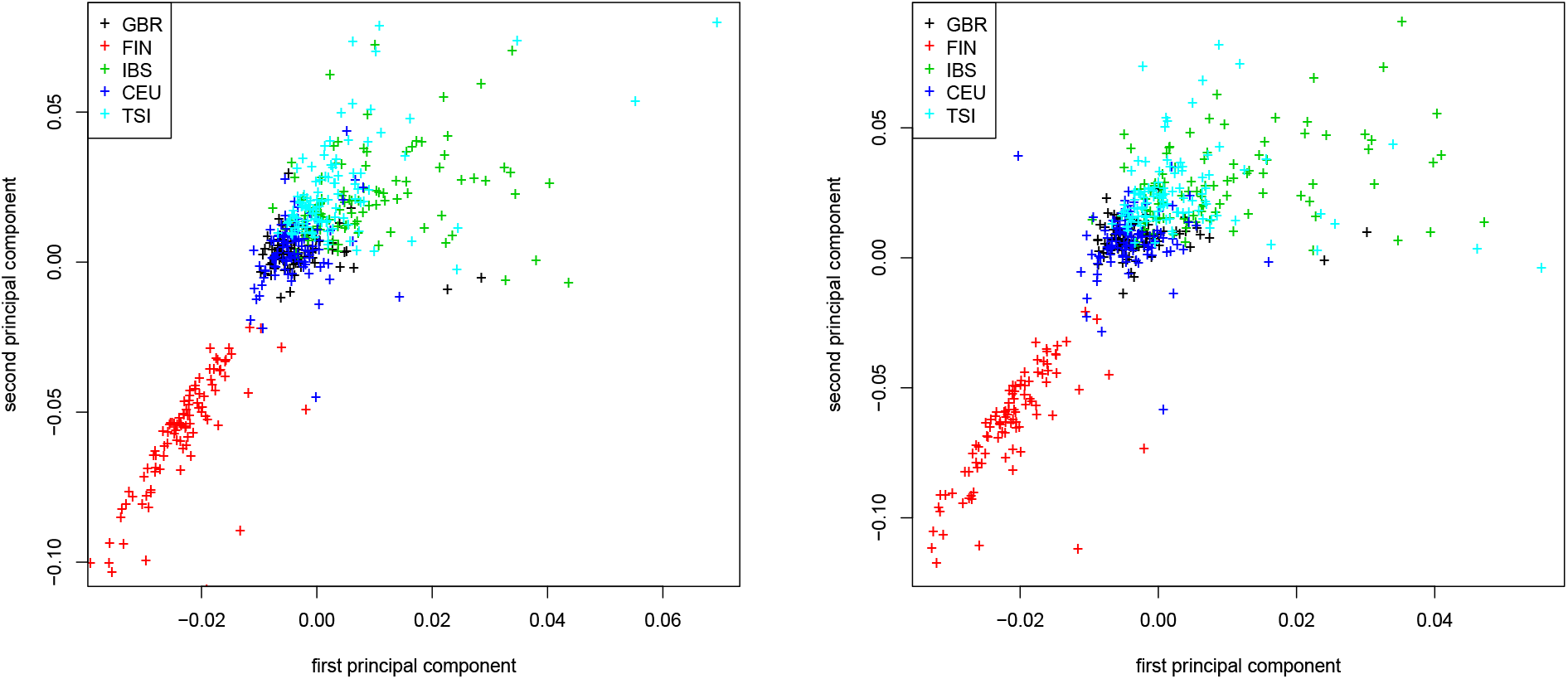
Genomic relationship matrix (GRM). First two principal components colored by population for the 1000 Genomes Project dataset. Traditional computation (left) and Algorithm 1 (right).

Figure 2 shows the first two eigenvectors of the original Jaccard and the approximate Jaccard matrix. We observe that here, the stratification plots are visibly different, though the approximate Jaccard matrix provides a very good stratification of the 1000 Genomes Project dataset.

**Figure 2:**
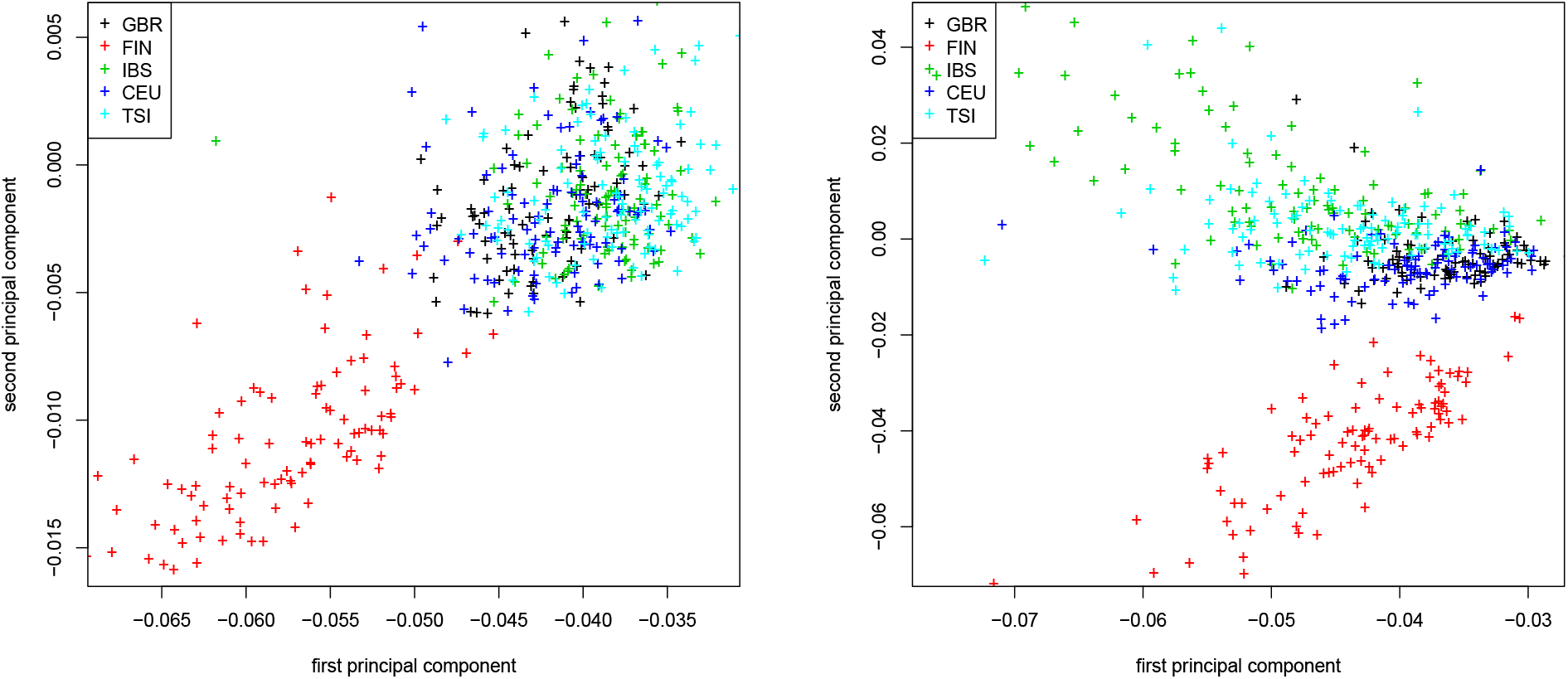
Jaccard matrix. First two principal components colored by population for the 1000 Genomes Project dataset. Traditional computation (left) and Algorithm 1 (right).

### 3.2 Verification of the theoretical bounds for the approximate Jaccard matrix

We are interested in quantifying further the numerical tradeoff made when computing the eigenvectors of the approximate Jaccard matrix. Additionally, we aim to verify the theoretical bounds on the approximate eigenvectors derived in Section 2.4.

To this end, we investigate both the validity and the scaling behavior of the bounds of Section 2.4 as a function of the number of variants *n*, the number of subjects *m*, the proportion of nonzero alleles, and higher order eigenvectors. We compute the Jaccard and approximate Jaccard matrices on simulated data. We create sparse matrices *G* of dimensions *n* × *m*, where a proportion *π* ∈ [0, 1] of entries is set to one (acting as nonzero alleles).

Figure 3 (left) shows results as the number of variants *n* is increased while keeping the number of subjects *m* = 100 fixed. The figure displays the measured *L*_2_ distance between the first eigenvector of the Jaccard matrix and the approximate Jaccard matrix, the angle bound of eq. (3), and the matrix bound of eq. (4). Similarly, in Figure 3 (right) we vary the number of subjects *m* while keeping the number of variants *n* = 100 fixed. In both cases, we observe that the measured *L*_2_ distance between the first eigenvector of the Jaccard matrix and its approximation is negligible. The angle bound (which requires the computation of both eigenvectors) seems to be a very tight bound, while the matrix bound (which requires no computation of eigenvectors and is thus an a priori bound) is valid but less tight. This is to be expected, as less information is required to compute the angle bound. In particular, in can be computed without having computed any eigenvectors.

**Figure 3:**
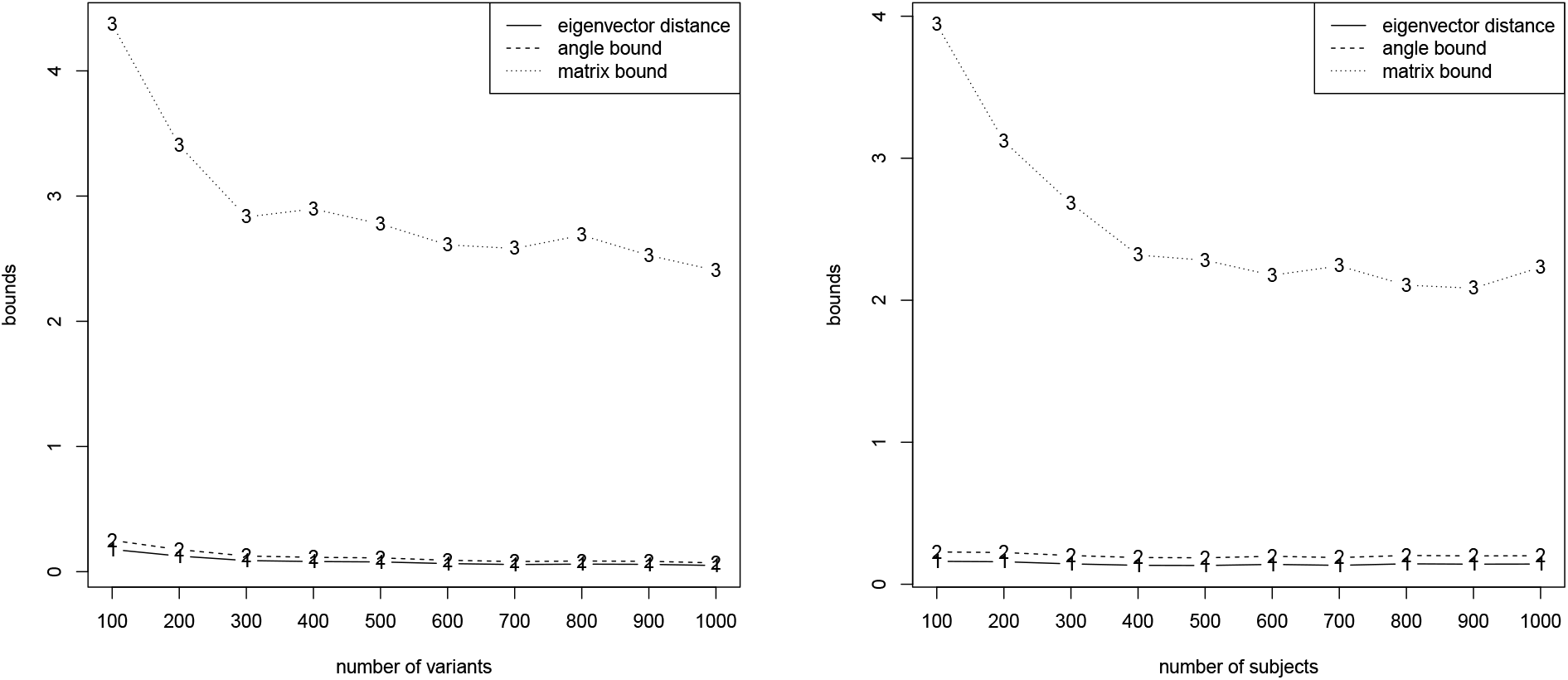
Measured *L*_2_ distance between the first eigenvector of the Jaccard matrix and the approximate Jaccard matrix, angle bound of eq. (3), and matrix bound of eq. (4). Variable number of variants *n* while keeping *m* = 100 fixed (left) and variable number of subjects *m* while keeping *n* = 100 fixed (right).

Figure 4 investigates the accuracy of the theoretical bounds as a function of the proportion *π* ∈ {0.1,…, 0.9} determining the sparseness of the matrix *G* while keeping the number of variants *n* = 1000 and the number of subjects *m* = 100 fixed. We observe that again, the error in *L*_2_ distance between the eigenvector computed for the Jaccard and approximate Jaccard matrices is negligible. The angle bound is again tight. The matrix bound is valid, more relaxed than the angle bound, and exhibits its closest bound for around *π* = 0.4.

**Figure 4:**
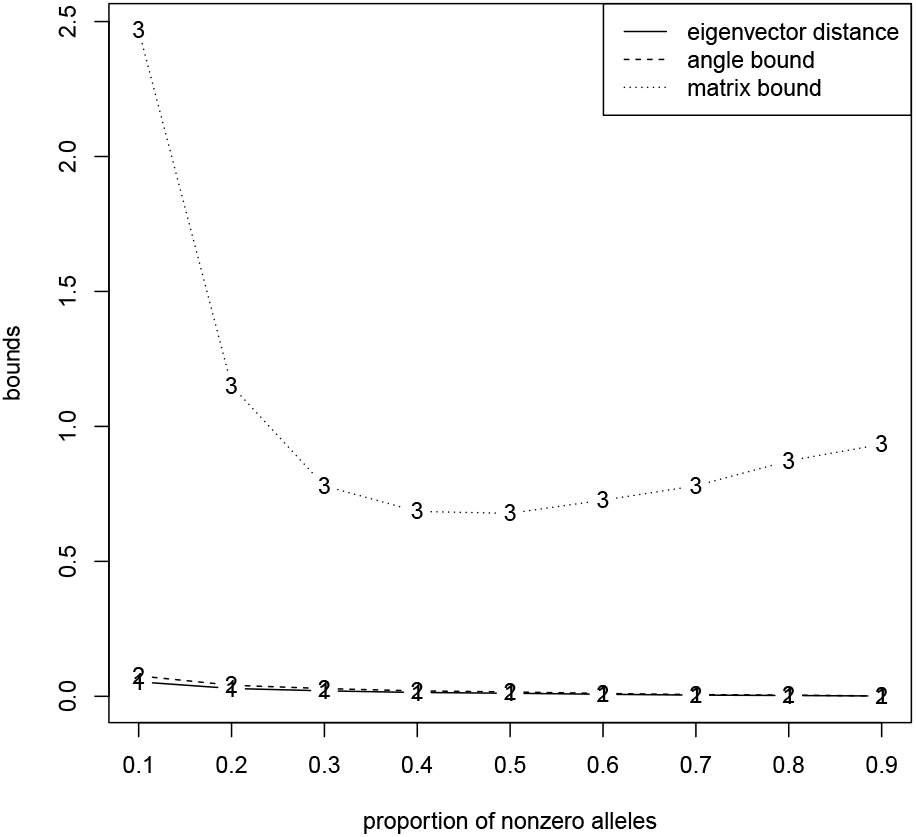
Measured *L*_2_ distance between the first eigenvector of the Jaccard matrix and the approximate Jaccard matrix, angle bound of eq. (3), and matrix bound of eq. (4). Proportion *π* ∈ {0.1,…, 0.9} of entries 1 while keeping the number of variants *n* = 1000 and the number of subjects *m* = 100 fixed.

Finally, Figure 5 applies the bounds of eq. (3) and eq. (4) to higher order eigenvectors. We observe that the approximate Jaccard matrix is more accurate for the first eigenvectors than the later ones. The angle bound nicely follows the actual observed *L*_2_ norm between the eigenvectors computed for the Jaccard matrix and its approximation, while the matrix bound is the same for all since it only takes the Jaccard matrix and its approximation into account (but no information on the eigenvector being computed).

**Figure 5:**
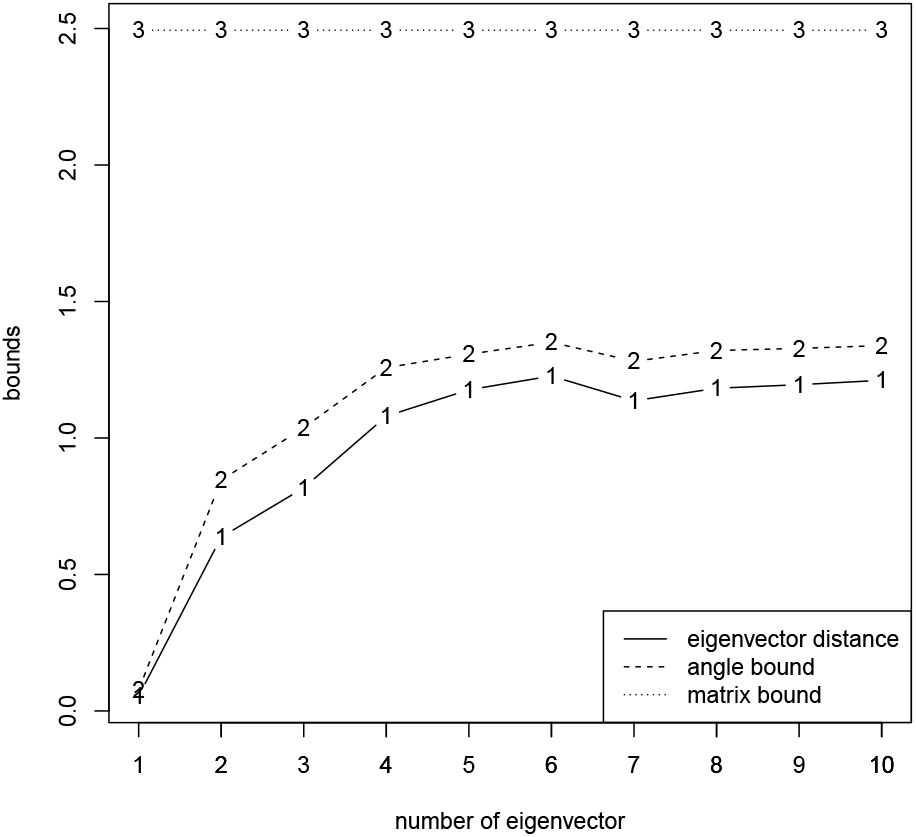
Measured *L*_2_ distance between the first eigenvector of the Jaccard matrix and the approximate Jaccard matrix, angle bound of eq. (3), and matrix bound of eq. (4). Bound progression for the first 10 eigenvectors while keeping the number of variants *n* = 1000, the number of subjects *m* = 100, and *π* = 0.1 fixed.

## 4 Discussion

This article considered the fast and efficient computation of the first eigenvectors of four similarity matrices, the genetic covariance matrix, the Jaccard matrix, the weighted Jaccard matrix, and the genomic relationship matrix. The computation of such eigenvectors is a standard tool in computational genetics, and an of importance for correcting linear regressions, revealing population stratification, and many more application areas.

In this contribution, we first introduce a unified way to express the genomic covariance matrix, the weighted Jaccard matrix, and the genomic relationship matrix which allows one to efficiently compute their eigenvectors in sparse matrix algebra using an adaptation of a fast SVD algorithm of Halko et al. (2011). An exception is the Jaccard matrix, which does not have a structure applicable for this computation. Second, we thus introduce a new approximate Jaccard matrix to which the fast SVD computation is applicable. Third, we establish guaranteed theoretical bounds on the distance (in *L*_2_ norm and angle) between the principal components of the Jaccard matrix and the ones of our proposed approximation. These a priori bounds allow the user to gauge in advance the trade-off between obtained speedup and sacrificed accuracy when using the proposed approximate Jaccard matrix.

## A Additional figures

Figure 6 and Figure 7 show analogous results as Figure 1 for the genetic covariance matrix and the weighted Jaccard matrix, respectively. Similarly to Figure 1, both are computed on the dataset of the 1000 Genomes Project. As seen in both figures, the population stratification plots obtained by either computing the similarity matrix first and then its eigenvectors via SVD, or by running Algorithm 1 are virtually identical. This is to be expected as the decomposition of Section 2.2 for the genetic covariance matrix and the weighted Jaccard matrix are exact.

**Figure 6:**
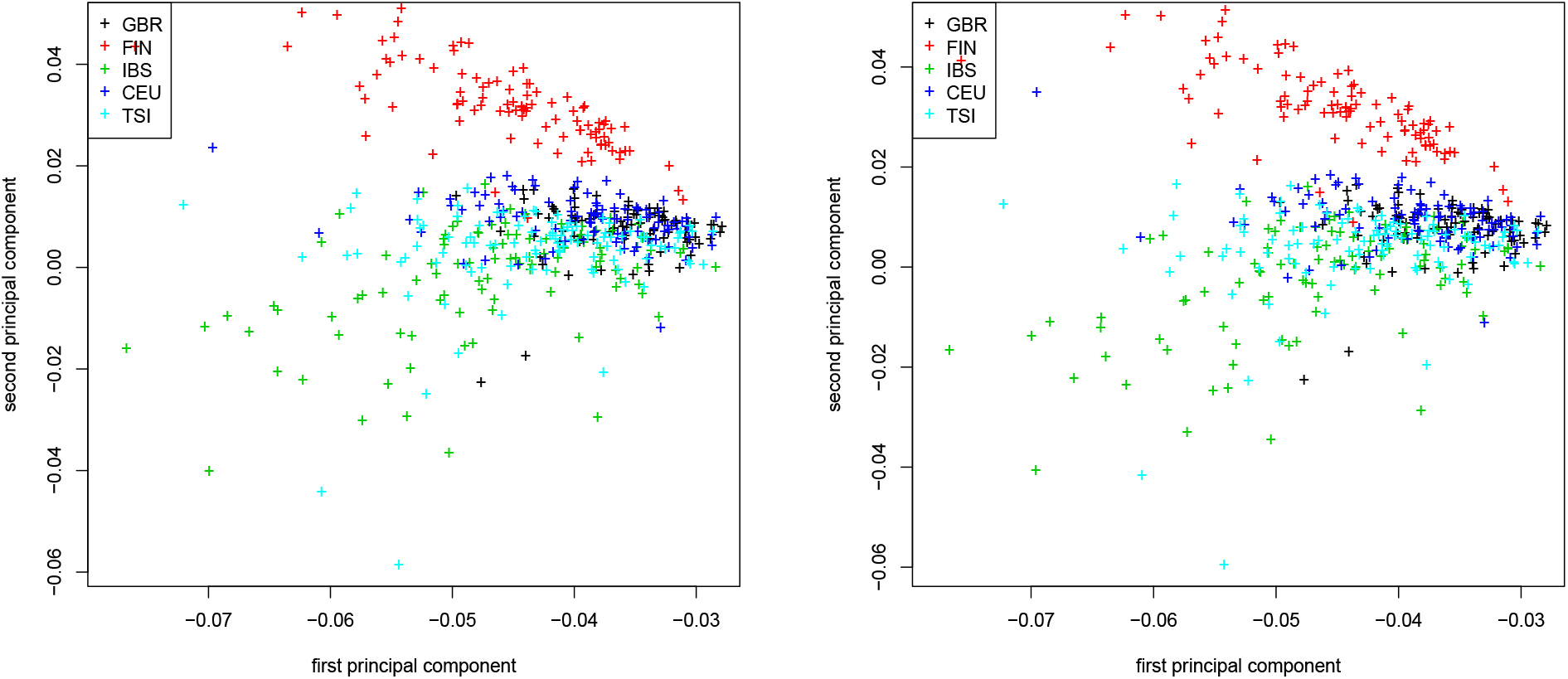
Genetic covariance matrix. First two principal components colored by population for the 1000 Genome Project dataset. Traditional computation (left) and Algorithm 1 (right).

**Figure 7:**
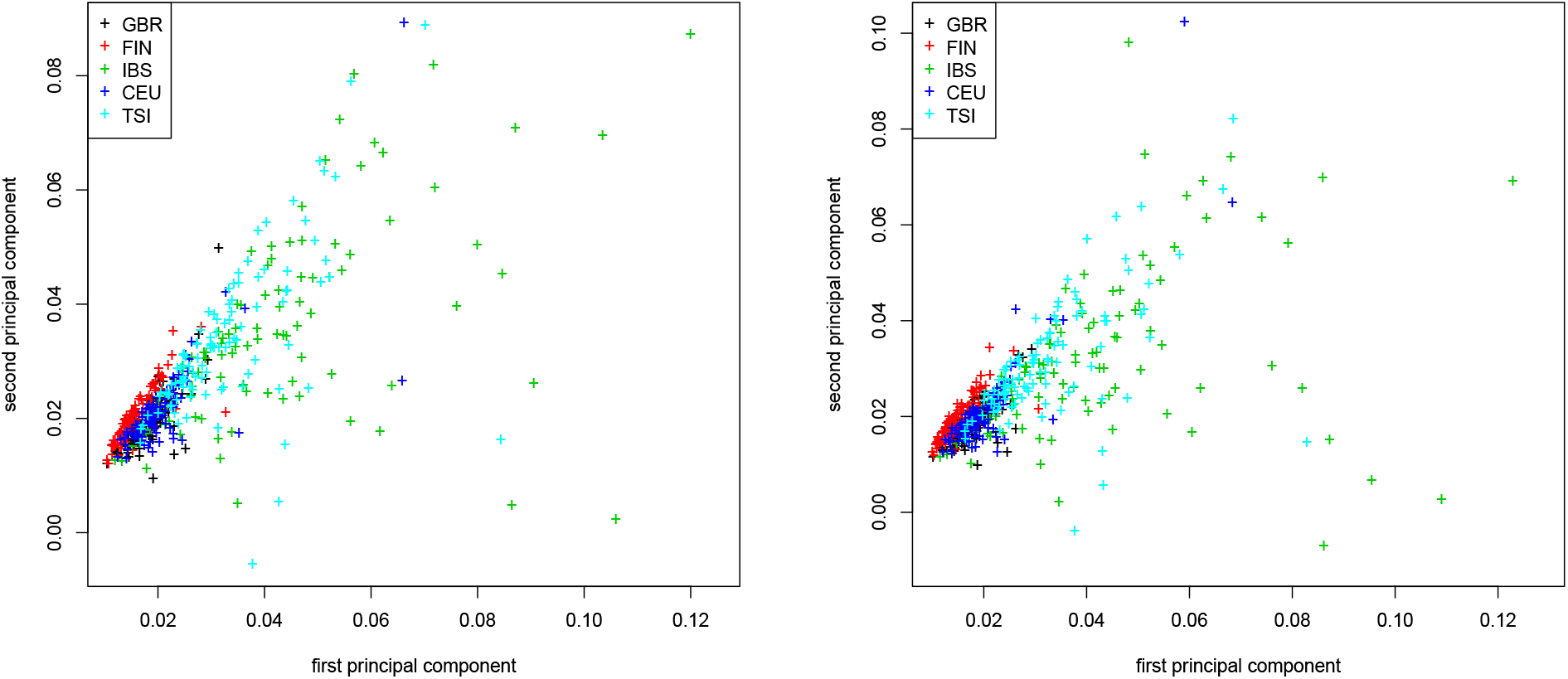
Weighted Jaccard matrix. First two principal components colored by population for the 1000 Genome Project dataset. Traditional computation (left) and Algorithm 1 (right).

